# Non-histone human chromatin protein, PC4 is critical for genomic integrity and negatively regulates autophagy

**DOI:** 10.1101/266932

**Authors:** Sweta Sikder, Sujata Kumari, Pallabi Mustafi, Nisha Ramdas, Swatishree Padhi, Arka Saha, Birendranath Banerjee, Ravi Manjithaya, Tapas K. Kundu

**Affiliations:** Transcription and Disease Laboratory, Molecular Biology and Genetics Unit, Jawaharlal Nehru Centre for Advanced Scientific Research, Bangalore, Karnataka, India; Molecular Stress and Stem Cell Biology Group, School of Biotechnology, KIIT University, Bhubaneswar, Odisha-751024, India; Autophagy Laboratory, Molecular Biology and Genetics Unit, Jawaharlal Nehru Centre for Advanced Scientific Research, Bangalore, Karnataka, India; Mechanobiology Institute & Department of Biological Sciences, National University of Singapore, Singapore; Transcription and Disease Laboratory, Molecular Biology and Genetics Unit, Jawaharlal Nehru Centre for Advanced Scientific Research, Bangalore, Karnataka, India.

**Keywords:** Nuclear architecture, chromatin organization, gamma irradiation resistance, epigenetic landscape, histone acetylation

## Abstract

Multifunctional human transcriptional positive co-activator 4 (PC4) is a bonafide non-histone component of the chromatin and plays a pivotal role in the process of chromatin compaction and functional genome organization. Knockdown of PC4 expression leads to a drastic open conformation of the chromatin and thereby altered nuclear architecture, defects in chromosome segregation and changed epigenetic landscape. Interestingly, these defects do not induce cellular death but result in enhanced cellular proliferation possibly through enhanced autophagic activity. PC4 depletion confers significant resistance to gamma irradiation. Exposure to gamma irradiation further induced autophagy in these cells. Inhibition of autophagy by small molecule inhibitors as well as by silencing of a critical autophagy gene drastically reduces the survivability PC4 knockdown cells. On the contrary, complementation with wild type PC4 could reverse this phenomenon, confirming the process of autophagy as the key mechanism for radiation resistance in absence of PC4. These data connect the unexplored role of chromatin architecture in regulating autophagy during a stress condition such as radiation.

## Introduction

Inside the limited space of the eukaryotic cell nucleus, the genome is packaged into a highly dynamic nucleoprotein structure called the chromatin, which regulates and supports critical cellular processes. In a given chromosome territory, the fine tuning of the three-dimensional homeostatic organization of the genome with proper epigenetic landscape is maintained by the non-coding RNA and several highly abundant non-histone chromatin associated proteins. The non-histone chromatin associated proteins which include high-mobility group proteins (HMGs)^1–4^, heterochromatin binding protein 1 (HP1)^5^, methyl CpG binding protein 2 (MeCP2)^6^, poly-ADP-ribose polymerase 1 (PARP-1)^7^, positive co-activator (PC4)^8^, among others, modulate the dynamicity of the genome organization by directly interacting with the different components of the chromatin including linker histones. Although functions of non-histone proteins are well known in context of gene regulation and expression, their intrinsic and detailed role as responders of environmental stress and thereby maintenance of cellular homeostasis remains to be understood in the global context.

Multifunctional human transcriptional coactivator, positive coactivator 4 (PC4) which was initially isolated from nuclear extracts of murine plasmacytoma^9^, was discovered as transcriptional co-activator known to facilitate activator dependent transcription by RNA polymerase II, *in vitro*, by directly interacting with upstream activators as well as the general transcriptional machinery.^10–11^ Sub1/PC4 is not only restricted to PolII driven transcription but is more diverse as it plays multiple roles in PolIII driven transcription^12^, DNA repair^13^ and replication.^14^ Previously, we have shown that PC4 is a bonafide non-histone component of the chromatin and induces its compaction by directly interacting with the core histones^8^. Transient silencing of PC4 led to an open chromatin conformation, altered epigenetic state and the expression of neuronal genes in non-neuronal cells.^15^ In one of the very recent reports we have shown that knockout of PC4 is embryonic lethal, however brain specific knockout leads to reduced neurogenesis with defects in memory extinction. The same study revealed that PC4 might play an important role in the regulation of hypoxia/anoxia and apoptosis or cell death.^16^

The human genome is efficiently safeguarded against environmental stresses and other DNA damage inducing agents by the coordinated functions of chromatin associated proteins (acting as sensors), altered histone modifications and thereby a changed transcriptional output.^17^ Macro-autophagy (herein autophagy), a well conserved cellular self-eating process, is a lysosome dependent degradative pathway known to maintain the cellular homeostasis by getting rid of unwanted organelles and toxic proteins. Autophagy has a prosurvival role as it responds to stress such as starvation and genomic insult due to radiation. Several studies have highlighted the cytoprotective role of autophagy upon radiation stress in tumour cells.^18^ This role might be mediated by extensive crosstalk of the autophagy process and the apoptotic pathway. It has also been shown that autophagy protects tumour cells from injury and stressful environment by eliminating unwanted or non-functional organelles, controlling the production of reactive oxygen species and by the recycling of essential proteins required for repairing DNA lesions.^18^ However, studies also reveal that radiation induced autophagy can have dual roles of cytoprotection as well as killing of cells. A study in glioma stem cells upon exposure to ionizing radiation led to induction of autophagy which helped in maintaining cellular viability. Treatment with small molecule inhibitors like Bafilomycin A1 and upon silencing of autophagy related genes like ATG5 and BECN1, these cells became sensitized to gamma irradiation which led to a significant decrease in the cell number.^19^ However, the phenomenon might not be universal as a more recent study showed that Rapamycin induced differentiation of glioblastoma multiform (GBM)-initiating cells (GICs) more sensitive to gamma irradiation by activating autophagy.^20^ Thus autophagy can act as a “double edged sword” conferring quite contrasting properties to the cells depending upon other extrinsic factors like the tissue specificity or the proteins regulating the induction of autophagy upon radiation.

The mechanism of autophagy under different stress stimuli is yet to be elucidated. There have been evidences that epigenetic modifications and thereby the chromatin state plays a critical role in altering the expression of key regulators of autophagy. Chromatin modulators like acetyltransferase, KAT5/TIP60 can be activated by GSK3 (glycogen synthase kinase-3), which in turn directly acetylates ULK1 and induces autophagy.^21^ Also the master lysine acetyltransferase p300 is known to negatively regulate autophagy, by its direct interaction with Atg7 in the cytoplasm.^22^ Knocking down of EP300 activates autophagy, whereas overexpression of EP300 inhibits starvation-induced autophagy. BAG6/BAT3 (BCL2-associated athanogene) was demonstrated to control basal and starvation-induced autophagy via limiting EP300-dependent acetylation of Atg7. There has been evidence that mammalian cells can degrade nuclear components by the process of autophagy to maintain the nuclear function and integrity.^23^ Autophagy is known to play a cytoprotective role under conditions of DNA damage, by regulating the dNTP pool levels,^24^ which are essential for DNA replication and repair. Thus, autophagy induction and initiation by various stressful environmental conditions is quite unique and is intricately regulated by the chromatin state, nuclear and cellular composition and architecture along with various modifications which the chromatin harbours. However, how these multiple factors act together in a concert and thereby balance the fate of death and survival of the cells in the context of autophagy remains to be found.

In the present study, we generated a stable PC4 knockdown cell line to explore the significance of PC4 in the physiological context. The PC4 depleted cells harbour an open or decompacted chromatin, and a significantly altered epigenetic landscape with severe morphological as well as segregation defects. In spite of acquiring such cellular defects, the PC4 deficient cells not only proliferated faster but also became more resistant to gamma irradiation-induced cytotoxicity. Significantly, the PC4 knockdown cells show greater induction of autophagy which further gets enhanced upon exposure to gamma irradiation. The gene network underlying the induction of this process of autophagy in absence of PC4 has also been investigated and it is shown to be mitigated through the global alteration of the epigenetic landscape. Our results highlight the critical role of autophagy in survival and radiation resistance of these PC4 depleted cells.

## Results

### Knockdown of PC4 leads to altered nuclear and chromosomal morphology and results in cell segregation defects

PC4, initially discovered as a transcriptional coactivator, was found to be a bonafide component of chromatin and it induced chromatin compaction both *in vitro* and *in vivo*.^8^ To explore the significance of PC4 in the physiological context, we generated a stable PC4 knockdown in HEK293 cell line (sh-PC4) using a lentivirus mediated shRNA delivery system (Fig. S1). Different shRNAs (marked as #) targeting different regions of PC4 gene were used to check the level of silencing of PC4 both at transcript and protein levels (Fig. S1A and B). The shRNA #5 was used to make a stable PC4 knockdown cell line as it showed maximum downregulation both at transcript and protein levels (Fig. S1A, B and C). In agreement with the transient silencing reported previously^15^, the stably transfected (#5 shRNA) PC4 knockdown cells showed up-regulation of GAD1, SCN2 and M4 gene expression (these are neuronal genes which are repressed by PC4) as compared to the control cells (Fig. S1D and E). Here in this study we have characterized in details the phenotype of PC4 knockdown cell line (harbouring shRNA #5) denoted as sh-PC4. The control cells used here is the wild type HEK293 cells. Although to negate the non-specific effect of shRNAs we established a control cell line (HEK293) stably expressing the scrambled shRNA clone (data not shown), but here all the assays were performed with wild type HEK293 as it replicated the same phenotype as seen in the non-silencing shRNA control cell line (data not shown). We observed that in absence of PC4, in the shRNA mediated knockdown cells, the individual metaphase chromosomes could not be spread unlike the chromosomes from the control cells. The typical metaphase chromosome shape was found to be altered in these cells (Fig. 1A). The shape and size of the nuclei was found to be highly variable and significantly deformed in the PC4 knockdown cells. Significantly the nuclear lamina also was invaginated when PC4 was absent in the nuclei, indicating the role of PC4 in the maintenance of the global nuclear architecture (Fig. 1B). The sh-PC4 cells also showed formation of anaphase bridges when binucleate analysis was performed upon nuclear staining with DAPI. (Fig. 1C). Anaphase bridge formation leads to segregation defects. Cells where PC4 was knockdown stably exhibited several signatures of defective segregation upon cell division such as unequal and irregular shaped daughter nuclei upon cell division with an increase in the number of anaphase bridges (Fig. 1D). The control cells however showed no such defects in the form of nuclear shape and size, and anaphase bridge formation also could not be detected. To visualize the segregation defects in greater detail at the chromosome level, telomere PNA-FISH (Peptide nucleic acid fluorescence *in situ* hybridization) was performed. In agreement with the anaphase bridge analysis, we observed binucleates with equal number of PNA foci as green signal in the control cells whereas in sh-PC4 cells, unequal number of PNA foci was predominant in the binucleates (Fig. 1E). Quantitation of the PNA signal showed that the unequal distribution of the foci in PC4 knockdown cells is statistically significant as compared to the near complete equal distribution in control cells (Fig. 1F). This unequal distribution of chromosomes in the newly divided daughter nuclei could be attributed to segregation defects. To visualize these structural anomalies in the PC4 depleted cells in a greater detail, both HEK293 and sh-PC4 HEK293 cells were subjected to transmission electron microscopy. The images showed abnormal nuclei with altered shape and accumulation of large number of double membraned vesicles around the nucleus of PC4 knockdown cells (Fig. 1G) which were completely absent in the control cells. Collectively, these results indicate that depletion of PC4 alters the nuclear as well as chromosomal morphology which in turn might also alter other cellular processes, thereby exhibiting an unexpected stressful cellular environment.

**Figure 1.**
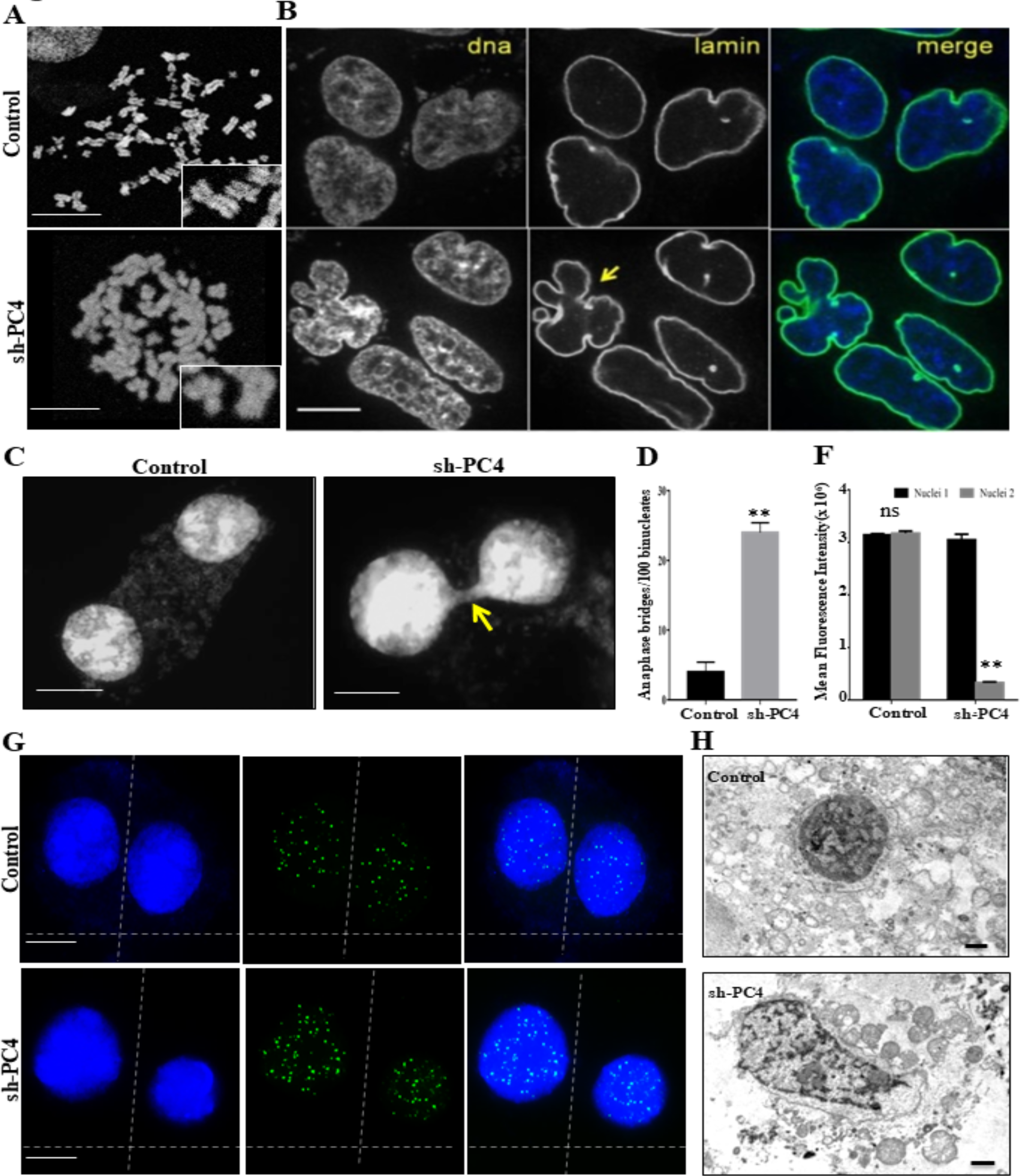
Knockdown of PC4 leads to altered nuclear and chromosomal morphology and defects in segregation. (A) Metaphase chromosome spreads reveal abnormal chromosomal structures in PC4 knockdown (sh-PC4) cells as compared to control cells. Scale bar 1μm. (B) Changes in nuclear shape and size with PC4 knockdown (sh-PC4) as compared to control cells, with top/bottom panels depicting changes to nuclear lamina (green) and DNA (blue) as indicated by arrows. Scale bar 2μm. (C) Analysis of nuclear division showing irregular segregation and formation of Anaphase Bridges in the sh-PC4 cells compared to control cells. Scale bar 1μm. (D) Quantitative representation of genome instability in terms of number of anaphase bridges formed per 100 nuclear divisions. Data are presented as means ± S.E.M. p** < 0.001, p*** < 0.0001. (E) Telomere PNA-FISH analysis showing irregular segregation in sh-PC4 cells in terms of differing number of PNA foci in daughter nuclei as compared to control. Green fluorescence represents Telomere-PNA probe. The horizontal dotted line denotes axis of seggregation while the vertical line denotes axis of cell division. Scale bar 1μm. (F) Quantitative representation of the PNA hybridization in daughter nuclei in terms of the mean fluorescence intensity in a.m.u. As shown, there is differing signal intensity in the daughter nuclei in sh-PC4 as compared to control cells. Data are presented as means ± S.E.M. (n = 3). p** < 0.001, p*** < 0.0001. (G) Representative image of electron micrographs from control versus sh-PC4 cells. sh-PC4 cells harbour nucleus of irregular shape and shows presence of vesicles near the nucleus. Scale bar 1μm.

### PC4 knockdown perturbs the epigenetic status

The ordered chromatin structure plays a vital role in maintaining the cellular homeostasis and is critical for efficient functioning of a cell. Considering the role of PC4 as a chromatin organiser, the compaction state of the chromatin upon knockdown of PC4 was analysed by digesting the nuclei of the knockdown cells with micrococcal nuclease in a time dependent manner. Compared to the control cells, the knockdown cells exhibit a decompacted chromatin as is evident by the presence of mono-nucleosomes in the digested pattern of MNase treated nuclei (Fig. 1A). When nuclei from both the cells were digested with equal units of MNase for similar time points as of 10 and 15 minutes, preponderance of mono-di-nucleosomes could be observed in the sh-PC4 cells as compared to the control cells (Fig. 2A, compare lanes 3-4 versus lanes 7-8). In control cells, at similar time points slow migrating higher molecular weight fragment could be observed as a smear thus signifying that the chromatin from the control cells were more resistant to MNAse digestion than the chromatin from sh-PC4 cells (Fig. 2A, lanes 3-4). It was also intriguing to observe that at 0 min time point of MNAse digestion (where the reaction was immediately stopped after addition of the enzyme), chromatin from sh-PC4 cells showed presence of mono-nucleosomes whereas the chromatin from the control cells showed no digestion pattern, as evident by the smear signifying higher molecular weight bands (Fig. 2A, compare lane 1 vs lane 5). This signifies the integral role of PC4 in maintaining the compacted chromatin state in cells. Thus, knockdown of PC4 not only alters the nuclear and cellular architecture, it also leads to an open chromatin structure. PC4 acts as a critical chromatin condensing protein, whose depletion in the cells leads to a remarkable change in the chromatin architecture. Subsequently, owing to the opening of the chromatin in the PC4 knockdown cells, we investigated the alteration in the epigenetic landscape especially in the context of histone modifications. All the transcriptional activation associated histone modification marks analysed, namely histone acetylation marks like H3K9Ac, H3K27Ac, etc. were found to be enhanced in the sh-PC4 cells as compared to the control cells (Fig 2B, compare lanes 1 vs 2). Interestingly, we also find upregulation of H3R17me2 mark which is in lieu with a previous report, of CARM1 regulating autophagy^25^. Taken together, these data establish that stable knockdown of PC4 not only alter the nuclear architecture but also the global chromatin organization towards a more open, and possibly transcriptionally amenable genome.

**Figure 2.**
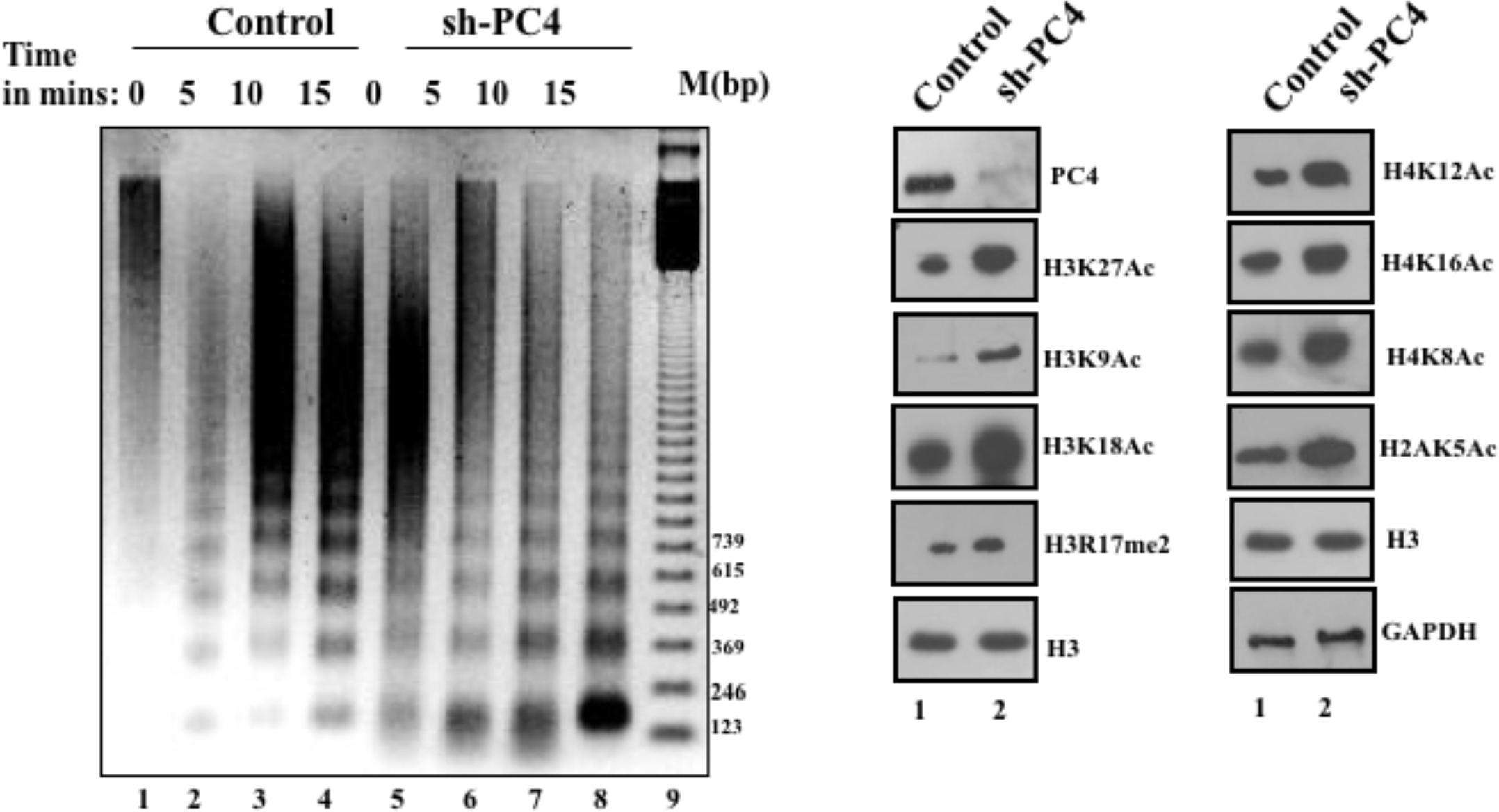
Absence of PC4 causes global chromatin decondensation, and consequent changes in epigenetic state. (A) Micrococcal nuclease digestions were carried out at four different time points (0, 5, 10 and 15 mins) with the nuclei isolated from control cells (lanes 1 to 4) and from sh-PC4 cells (lanes 5 to 8). (B) Comparison of histone modification marks between control and sh-PC4 cells. (C) Core histones extracted from control cells and knockdown cells. Transcription activation associated histone modifications were examined by immunoblotting using highly specific antibodies against specific histone modifications as indicated.

### Knockdown of PC4 exhibits higher cell migration and enhanced survival upon exposure to gamma irradiation

The PC4 knockdown cells were examined further for their sustainability through cell division. Surprisingly we found that these cells were able to divide and grow well in multiple passages; infact knockdown cells appeared to grow faster than control cells. To understand the rate of proliferation upon PC4 knockdown, we performed wound healing assays by creating equal sized scratch by a sterile tip. Healing was much faster in case of sh-PC4 when compared to control cells (Fig. 3A). The migratory ability of cells was also compared upon passing high current through the cells. After 12 hours, impedance of sh-PC4 cells was still higher than the control cells indicating PC4 knockdown cells have higher migration ability (Fig. 3B). The faster healing of wound could also be attributed to higher proliferation rate of sh-PC4 apart from its greater cell migration ability than control cells. The sh-PC4 cells indeed showed higher proliferation rate as is evident by the clonogenic assay (Fig. 3C). On seeding equal number of cells for both sh-PC4and control cells, PC4 depleted cells not only showed greater number of colonies after 10 days of growth as compared to the control cells but the colonies also appeared to be larger in size and shape, morphologically. However, increased proliferation despite the several defects in sh-PC4 cell nuclei was puzzling. To get an insight into the mechanism of the higher proliferation ability of sh-PC4 cells, we further exposed these cells to a genotoxic stress like gamma irradiation and then compared the proliferation rates as well as cell death in control and PC4 knockdown cells. Remarkably, greater survival was observed in case of PC4 knockdown cells upon increased gamma radiation as compared to control cells as evidenced by appearance of significantly greater number of live colonies after 10 days of exposure to gamma irradiation (Fig. 3D and 3E). We also investigated the state of apoptosis in the sh-PC4 cells upon exposure to gamma irradiation by flow cytometry analysis using Annexin VCy3.18 dye. Annexin-Cy3.18 (AnnCy3) binds only to apoptotic cells which can be detected as red fluorescence which has been represented as blue in the upper quadrant (Fig. 3F). The live cells were pseudocolored as red. sh-PC4 cells harbour the stably expressing GFP containing shRNA plasmid (pGIPZ-shRNA for PC4 refer to materials and methods section). The stably transfected sh-PC4 cells thus emitted GFP fluorescence detected as green signal, in the lower quadrant of the sh-PC4 cells (Fig. 3F). Using Annexin V staining coupled to flow cytometry, we observed that sh-PC4 cells apoptosed significantly lesser than the control cells upon exposure to gamma irradiation even after 6hours of exposure (Fig. 3F). The control cells showed a significant increase in the percentage of apoptotic cells 6 hours after exposure to gamma irradiation (4.76% to 11.43%) while the percentage of apoptotic cells in sh-PC4 cell population showed no significant alteration in their population even after exposure to gamma irradiation (Fig. 3G). These results signify that PC4 depletion confers radiation resistance to the cells.

**Figure 3.**
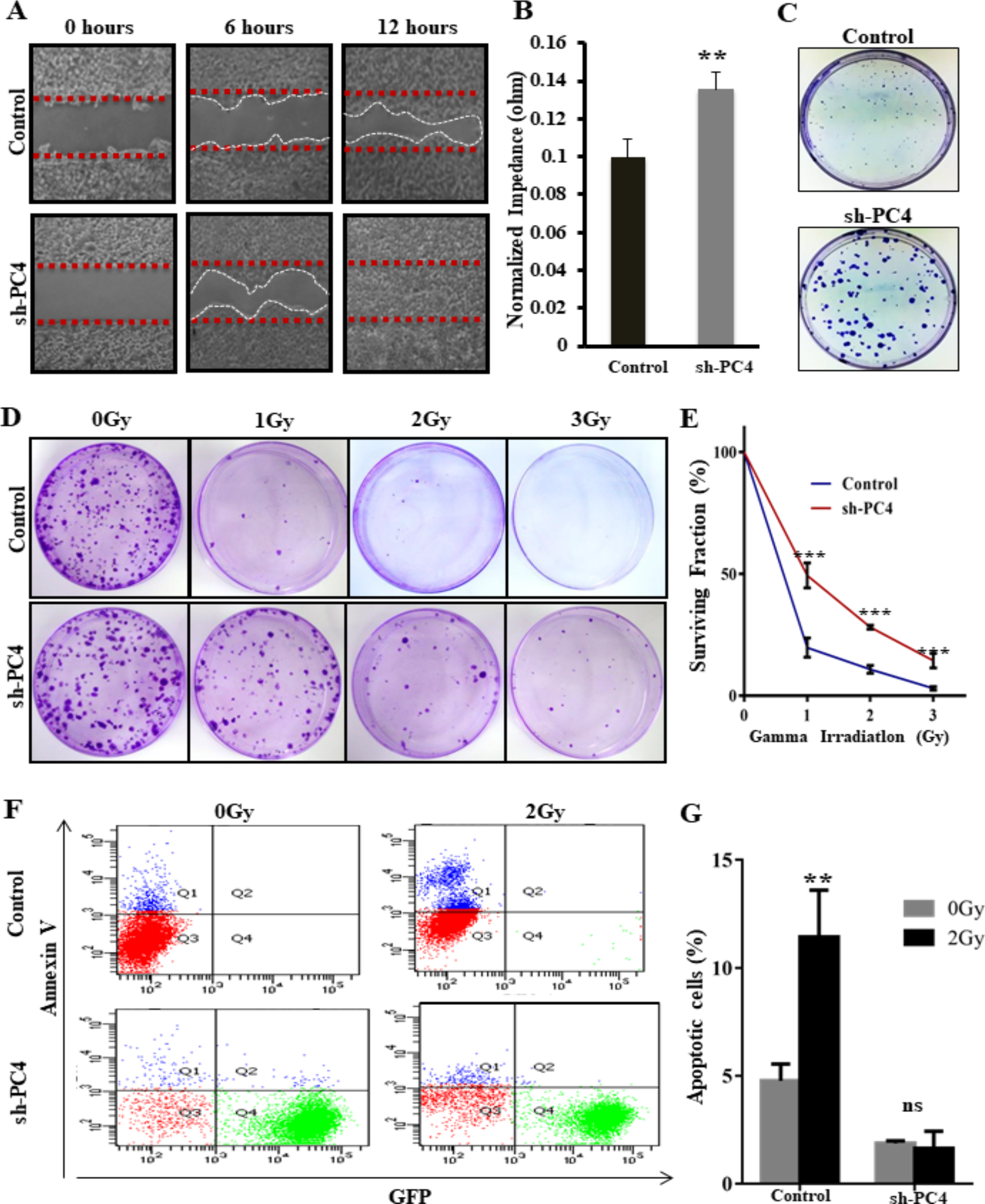
Absence of PC4 renders radioresistance upon exposure to gamma irradiation. (A Microscopic images of wound healing assay performed towards comparison of proliferation rate between control and sh-PC4 cells. Equal wound created in both cell lines (0 hours) was monitored at regular time intervals till complete healing was achieved in either of the cells (12 hours). (B) The representation of the normalized impedance for the control versus sh-PC4 cell population healing the wound for the monitored period using real time cell analyser. Data are presented as means ± S.D. (n = 3). p** < 0.001, p*** < 0.0001 (C) Representative images of the crystal violet stained colonies of control and sh5 cells shows greater proliferation rate in sh-PC4 cells. (D) Representative images of the crystal violet stained colonies of control and sh-PC4 cells sustained after exposure to increasing doses of the gamma radiation. (E) Graph plot depicting surviving fraction in percentage. Data are presented as means ± S.E.M. (n = 3). p** < 0.001, p*** < 0.0001. (F) Apoptosis analysis was performed in control vs sh-PC4 cells. Cells were irradiated at 2Gy and after 6hours of exposure; cells were analysed by FACS after staining the cells with Annexin V. The upper right and left quadrant represents apoptotic cells. Since sh-PC4cells harbour a GFP expressing shRNA plasmid, the upper left quadrant in sh5 panel represents apoptotic GFP positive cells. (G) Total percentage of apoptotic cells were counted from the FACS analysis and represented in the bag graph. Data are presented as means ± S.E.M. (n = 3). p** < 0.001, p*** < 0.0001.

### PC4 knockdown cells exhibit enhanced autophagy levels

In order to elucidate the molecular mechanism of the unique ability of sh-PC4 cells to survive better inspite of harbouring severe cell segregation and nuclear defects, we hypothesized that a compensatory cellular pathway such as autophagy might be operating rendering the property of enhanced proliferation and resistant to gamma irradiation to these cells. Autophagy, a well conserved, cellular degradative process is known to act as a potent survival mechanism for cells especially under stress or toxic conditions.^26^ We thus looked into the autophagy levels in sh-PC4 cells, by monitoring the levels of microtubule-associated protein light chain 3B (LC3) protein (an autophagy marker). LC3 is known to undergo lipidation upon induction of autophagy, the lipidated LC3 (LC3II) can be monitored in a SDS-PAGE as it acquires differential mobility as compared to the non lipidated form (LC3I).^27^ The western blotting analysis of cell lysates from PC4 knockdown and control cells show that in sh-PC4 cells the levels of LC3II (lipidated form of LC3) was significantly enhanced as compared to the control cells (Fig. 4A, compare lane 1 versus 2). Furthermore, when cells were grown in a nutrient deficient media for 2 hours to induce autophagy, sh-PC4 cells showed further increase in the levels of LC3II (Fig. 4A, lane 3 versus 4). The increase in the LC3II level can also be attributed to the accumulation of autophagy vacuoles due to a defective autophagy pathway; to investigate this possibility bafilomycin inhibition assay was carried out. Bafilomycin disrupts the autophagic flux by inhibiting the lysosomal proton pump V-ATPase, resulting in a defect in autophagosome-lysosome fusion.^28^ Upon treatment with bafilomycin (50nM and 100nM), there was a significant increase in the LC3II levels in the sh-PC4 cells over and above the induced LC3II levels (Fig. 4B, lane 4 versus 5 and 6). These data argue for the fact that the induced LC3II level observed in the sh-PC4 cells is indeed due to enhanced autophagy which is abrogated in the presence of an inhibitor, bafilomycin. To detect the cellular signatures of autophagy in the PC4 knockdown cells, both the control and PC4 knockdown cells were subjected to electron microscopy. In agreement with the biochemical data, the cellular visualization showed accumulation of large number of double membraned vesicles in the sh-PC4 cells which are characteristics of autophagy vesicles (Fig. 4C, lower panel). Visualization of such vesicles was not observed in the control cells even after imaging several fields of electron micrographs (Fig. 4C, upper panel). Presence of autophagy like double membraned vesicles even in the absence of autophagy inducers in the sh-PC4 cells further validate the fact that basal autophagy levels is elevated in the absence of PC4.

**Figure 4.**
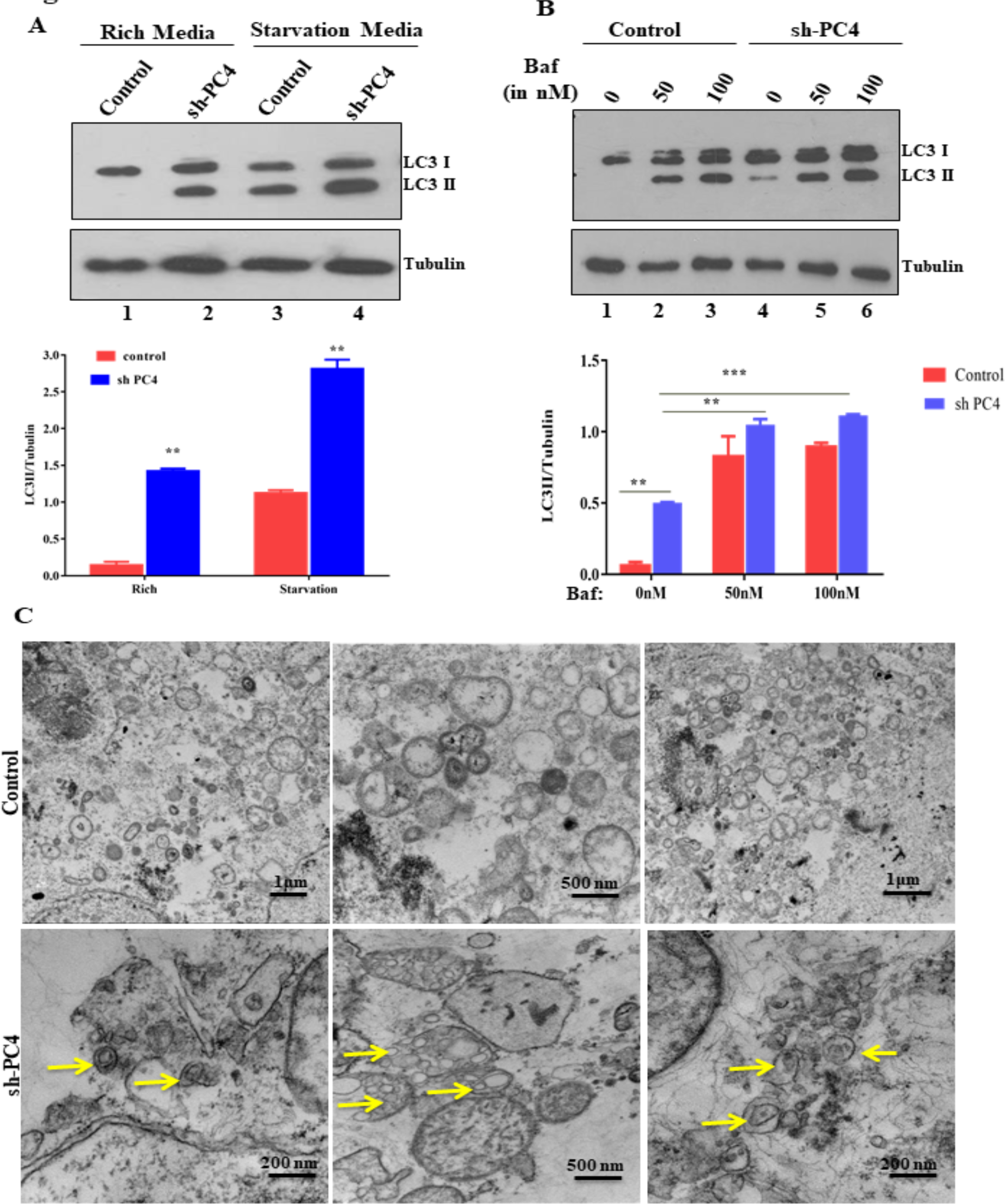
Absence of PC4 induces autophagy. (A) Induction of autophagy in sh-PC4 cells upon starvation: Control and sh-PC4 cells were starved for 2 hours in EBBS media. Cell lysates from both control and sh-PC4 cells in rich and starvation media were analyzed for LC3I and LC3II levels. Tubulin used as the loading control. Representative graph showing the change in LC3II level upon PC4 knockdown both in rich and starvation media is shown. Data are presented as means ± S.E.M. (n = 2). p* < 0.01, p** < 0.001, p*** < 0.0001 (B) Effect of Bafilomycin (50 nM, 100nM) on control and sh-PC4 cells. Accumulation of LC3II levels upon Bafilomycin treatment for 2 hours signify an inhibition in the process of autophagy. Corresponding, representative graph showing the change in LC3II level upon Bafilomycin inhibition both in control as well as sh-PC4 cells. Data are presented as means ± S.E.M. (n = 2). p* < 0.01, p** < 0.001, p*** < 0.0001 (C) Electron micrographs of sh-PC4 cells (lower panel) show huge accumulation of double membraned vesicles which are characteristics of autophagy vacuoles. No such vacuoles were observed in control cells (upper panel). Scale bar as denoted in figure.

### Induced autophagy in PC4 depleted cells harbours gamma radiation resistance to the PC4 knockdown cells

Since sh-PC4 cells show enhanced autophagy even without starvation, stress or any chemical autophagy enhancer, we wanted to investigate the functional role of autophagy in conferring radiation resistance to PC4 knockdown cells. For this purpose, both the control and sh-PC4 cells were exposed to increasing doses of gamma irradiation. Western blot analysis of cell lysates from sh-PC4 cells and control cells after 24 hours of gamma irradiation showed enhanced LC3II levels in sh-PC4 cells as compared to the control cells suggesting greater induction of autophagy in the PC4 depleted cells as compared to control cells (Fig. 5A, compare lanes 3 versus 4, 5 versus 6, and 7 versus 8, respectively). This observation suggests that gamma irradiation further triggers the autophagy levels in the PC4 depleted cells which might be playing a role in conferring its better survivability upon the cytotoxic cellular stress condition. To elucidate this hypothesis, we pre-treated the cells with two known small molecule autophagy inhibitors, 3-Methyladenine (3-MA) and Bafilomycin (Baf). 3-MA is known to inhibit autophagy by blocking autophagosome formation by inhibiting the type III Phosphatidylinositol 3-kinases (PI-3K), making it an early inhibitor of the autophagic pathway^28^. Pre-treatment with both 3-Methyladenine and Bafilomycin led to significant decrease in the number of live sh-PC4 cells as compared to the control cell line. (Fig. 5B and 5D). Administration of autophagy inhibitors also led to a significant decrease in the viability of sh-PC4 cells which were exposed to gamma irradiation (Fig. 5B and 5D). To further validate that autophagy is critical for enhanced survivability of sh-PC4 cells, we resorted to genetic inhibition of autophagy pathway by administering cells with specific shRNA against ULK1 gene. ULK1 is known to be act at the initiation of the autophagosome formation; its inhibition has been shown to shut down autophagy in several cell lines. Interestingly, we find that shRNA mediated knockdown of ULK1 drastically reduced the proliferation rate of the sh-PC4 cells in comparison to the control cells. (Fig. 5H and 5I). Collectively these data indicate that enhanced autophagy is one of the key mechanisms of better survivability and enhanced proliferation of PC4 depleted cells.

**Figure 5.**
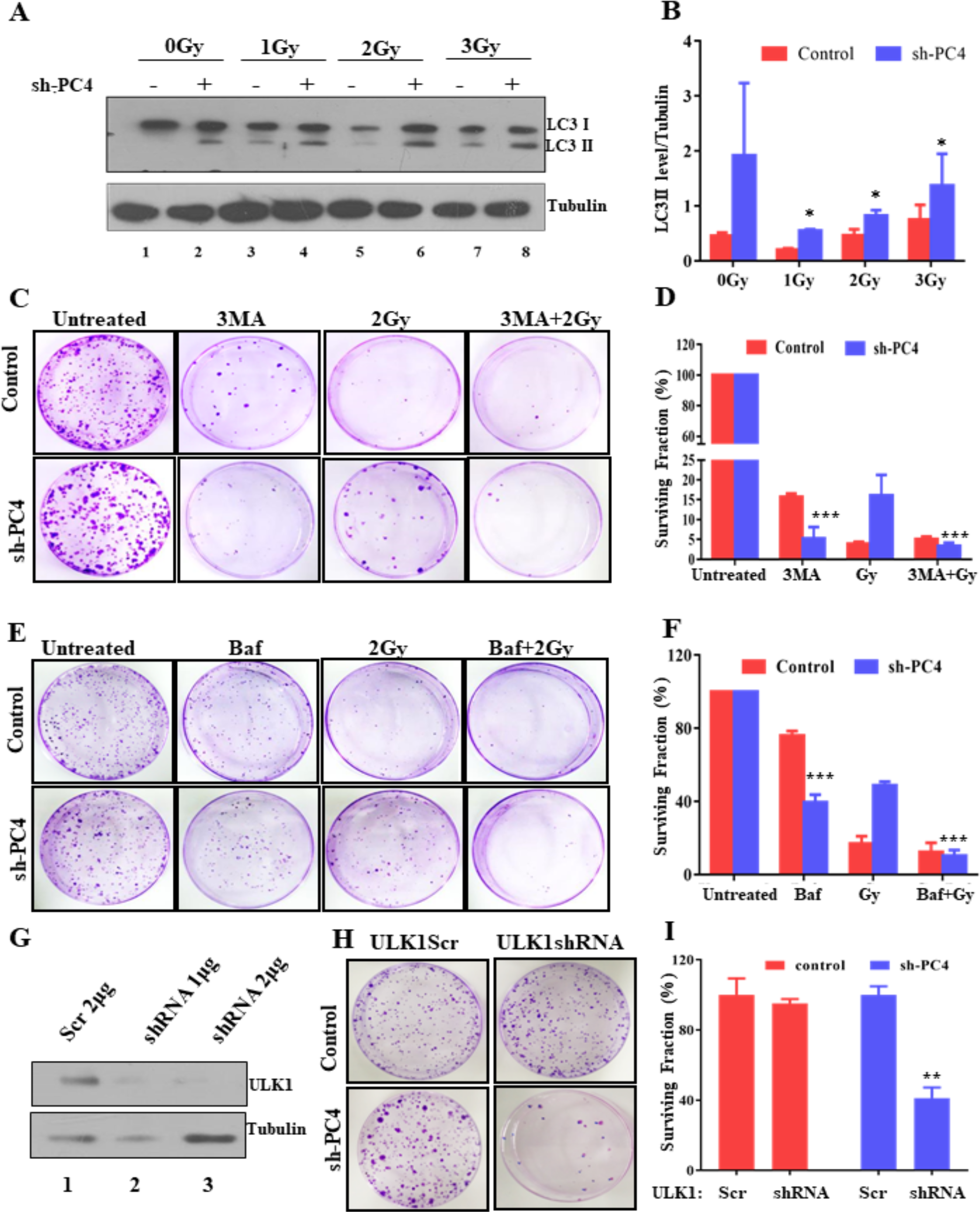
Exposure to gamma irradiation in absence of PC4 induces autophagy further. (A) Induction of autophagy in sh-PC4 cells upon gamma irradiation: Control and PC4 knockdown sh-PC4 cells were irradiated with different doses of gamma rays as indicated. Cell lysates from both cell lines were harvested after 24 hours post irradiation and western blot analysis were carried out with specific antibodies. LC3 antibody was used to analyse the process of autophagy while tubulin was used as loading control. (B) Representative graph showing the change in LC3II level upon gamma irradiation both in control and sh-PC4 cells is shown. Data are presented as means ± S.E.M. (n = 2). p* < 0.01, p** < 0.001, p*** < 0.0001. Autophagy inhibition sensitizes PC4 depleted cells to gamma irradiation: Control and sh-PC4 cells were treated with small molecule inhibitors of autophagy 10 mM 3-methyladenine (3-MA) (C) and 100nM Bafilomycin (Baf) (E) 2 hours prior to exposure to 2Gy of gamma irradiation. Cells were grown in the presence of the respective inhibitor for 24 hours post irradiation. Colonies were grown for 10 days. (C) and (E) Representative images of the crystal violet stained colonies of control and sh-PC4 cells sustained after treatment with 3MA and Bafilomycin and exposure to gamma irradiation. (D) and (F) Bar Graph depicting surviving fraction in percentage of the control and sh-PC4 cells after autophagy inhibition. Data are presented as means ± S. E. M. (n = 2). p* < 0.01, p** < 0.001, p*** < 0.0001. (G) Western blotting to confirm ULK1 knockdown 48 hours post transfection with specific shRNA against ULK1 in increasing concentrations. (H) Control and sh-PC4 cells were transfected with 2 μg of shRNA for 48 hours, and then plated for colony formation assay. Representative images of the crystal violet stained colonies of control and sh-PC4 cells sustained after transfection of ULK1shRNA. (I) Bar Graph depicting surviving fraction in percentage of the control and sh-PC4 cells after autophagy inhibition. Data are presented as means ± S.E.M. (n = 2). p* < 0.01, p** < 0.001, p*** < 0.0001.

### PC4 knockdown induces transcription of autophagy related genes in an epigenetic manner

Absence of PC4 creates open chromatin architecture in the nucleus with transcriptionally favourable epigenetic state of the genome. Therefore, the molecular mechanism of PC4-contextual autophagy enhancement could be causally related to the altered gene expression. To investigate whether depletion of PC4 has a direct transcriptional output of autophagy related genes we followed the possible gene expression signature in PC4 knockdown cells both before and after exposure to gamma irradiation. Control and sh-PC4 cells were exposed to 2Gy of gamma irradiation. After 6 hours of irradiation, the live cells were sorted using Annexin V cy3 FACS analysis. The live cell population was sorted out from the positively stained apoptotic cell population (Annexin V cy3.18 positive) from both the control and sh-PC4 cells before and after gamma irradiation. RNA extraction was carried out from the above mentioned live cell population from control as well as sh-PC4 cells, both before and after gamma irradiation. This RNA was further subjected to qPCR analysis to investigate the differential gene expression signature between the control and sh-PC4 cells upon gamma irradiation. Fig. 6A summarizes the result obtained for gene expression of each autophagy related genes as obtained from real time PCR analysis. Different colour represents different autophagy genes; expression of each gene is denoted by four bars, first represents expression in control cells, second fold change in gene expression in sh-PC4 cells, third represents the fold change in gene expression after 6 hours of gamma irradiation (2Gy) in control cells and fourth represents the same treatment but in sh-PC4 cells. The real-time PCR analysis reveals upregulation of most autophagy genes like AMPK, ULK1 and ULK2, MAP1LC3A and MAP1LC3B, etc., in sh-PC4 cells as compared to the control cells (Fig. 6A, compare first bar to second bar of each colour coded gene) which gets further upregulated upon exposure to gamma irradiation in sh-PC4 cells (Fig. 6A, compare third bar to fourth bar of each color coded gene). Notably in accordance with the protein levels (Fig. 5A lane 1 versus 3), the control cells showed induction of autophagy upon gamma irradiation also at the transcript level. This further verifies the fact that the induction of autophagy is a prime factor that might be responsible for mediating the cell survivability even for control cells as the gene expression analysis was carried out from the live population of control cells. For the knockdown cells, although we observe that there is a trend of upregulation of genes related to autophagy upon gamma irradiation it is not as significant as in case of control cells. This could be due to the significant enhancement of the expression of autophagy genes in PC4 knockdown cells even without the exposure of gamma irradiation which may fulfil the requirement of the gamma irradiation induced autophagy. Thus, PC4 knockdown elevates the transcription of the autophagy related genes leading to a highly activated autophagic pathway functioning in the sh-PC4 cells. Collectively, we observed that several genes related to the different steps of autophagy are upregulated upon PC4 knockdown at transcriptional level. To understand the selectivity of the phenomenon we also assayed the expression of certain genes unrelated to the autophagic pathway. Significantly it was observed that expression of these genes, namely Glyceraldehyde 3-phosphate dehydrogenase (GAPDH), histone acetyltransferase p300, Synaptonemal complex protein (SCP1) do not change upon PC4 knockdown in contrast to the genes related to autophagy pathway (Fig. S2D).

**Figure 6.**
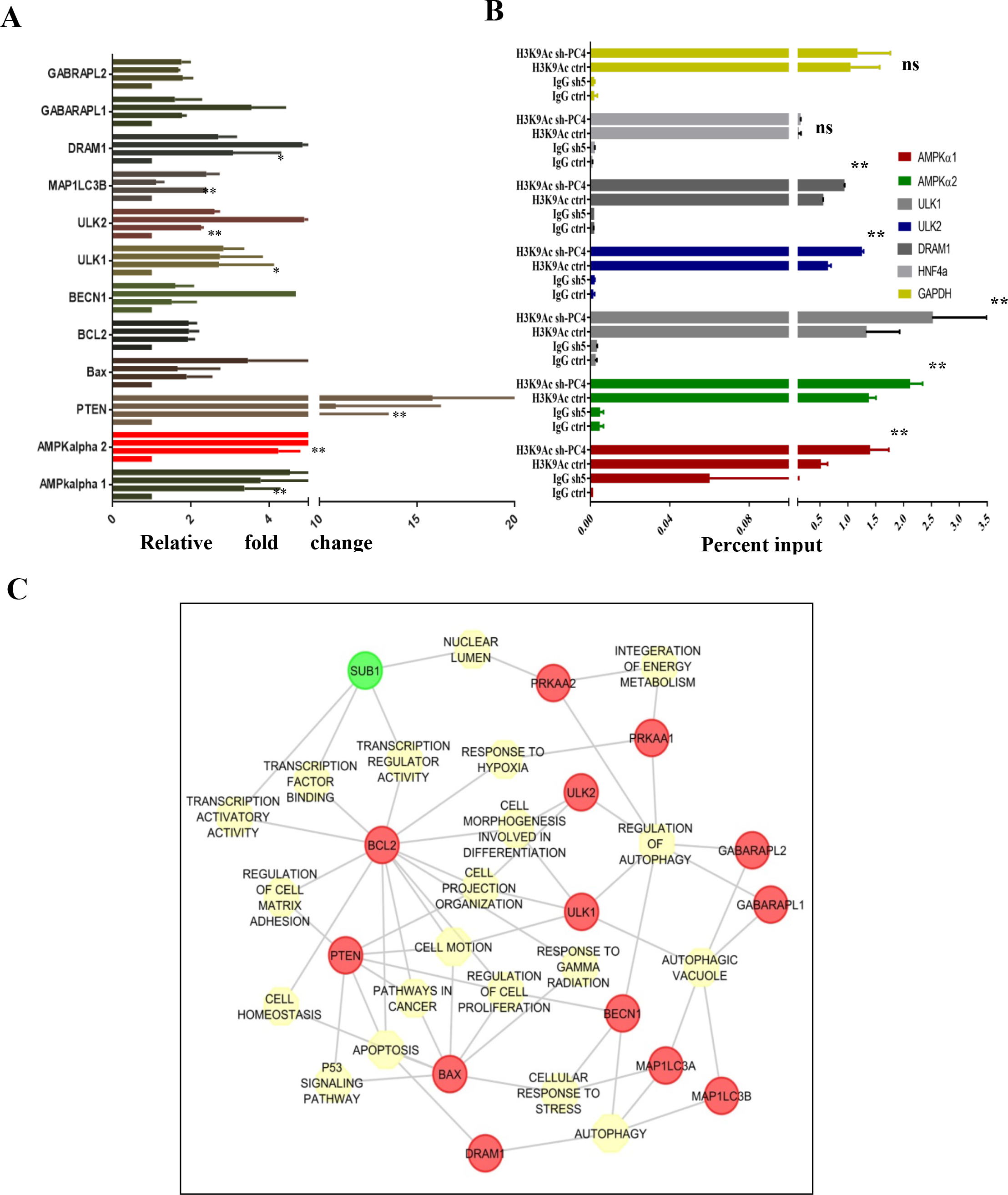
PC4 regulates autophagy related genes at the transcriptional level. (A) Alteration of autophagy related genes as determined by qPCR in ctrl versus sh-PC4 cells before and after gamma irradiation. Data are presented as means ± S.E.M. (n = 2) p** < 0.001, p*** < 0.0001. Actin is used as an internal control. (B) Occupancy of H3K9Ac at the promoters of selected autophagy related genes were assayed in both control and sh-PC4 cells. Percentage of enrichment of the mark at the selected promoters over the input was calculated. Data are presented as means ± S.E.M. (n = 2). p** < 0.001, p*** < 0.0001. (C) Altered gene network upon PC4 knockdown shows alteration in autophagy related genes and thereby the associated gene network. Down regulated gene is colour coded as green while genes showing upregulation is denoted as red.

Since absence of PC4 enhanced the histone acetylation related to transcriptional activation, we investigated the level of histone H3K9 acetylation (H3K9Ac) on the promoter sites of different upregulated genes by chromatin immunoprecipitation assay using highly specific antibodies for H3K9Ac. We investigated the occupancy of the active histone H3K9Ac marks at the promoters of 5 autophagy related genes which were known to be essential and critical in the initiation of the autophagic pathway in the cells. ChIP followed by qPCR with specific primers against the promoter region of each selected gene, yielded enhanced occupancy of the H3K9Ac mark at the promoters of the autophagy genes in sh-PC4 cells as compared to the control cells (Fig. 6B). The ChIP qPCR results are represented as the percent input; second bar for each individual colour coded gene represents enrichment in sh-PC4 cells while the first bar is representative of H3K9Ac occupancy in the control cells (Fig. 6B). In the promoters of all the 5 autophagy genes that we tested, the H3K9Ac occupancy was enhanced in the sh-PC4 cells as compared to the control cells. However, the occupancy of the active H3K9Ac mark do not alter at the promoters of genes like GAPDH and HNF4a (genes non-related to autophagy pathway) signifying that the occupancy of the specific active mark is somewhat specific in coherence with the transcriptional output result summarized before. These results are in agreement with the fact that H3K9Ac mark is highly upregulated in absence of PC4 in the cells which gets possibly enriched at specific promoter sites in the genome leading to an altered cellular physiology. Thus, PC4 regulates the autophagy related genes possibly at the transcriptional level by epigenetically modulating their promoters. To understand the global change in the genome architecture and thereby the transcriptional alteration in PC4 depleted conditions, we resorted to *in silico* analysis of pathways regulated by the different altered genes (protocol described in the materials and methods). This *in silico* network analysis of pathways interconnects several genes possibly regulated by PC4, and thereby providing an insight to the cellular pathways which might get altered in PC4 depleted cells. The analysis provides a broader overview of the altered physiological state of the PC4 depleted cells which might be due to the concerted effect of the alteration of the several pathways as denoted on the line interconnecting the nodes. Most of the genes to be altered belonged to the autophagy regulatory pathway referred in red circular nodes (Fig. 6C). It was also interesting to observe that perturbation of PC4 levels not only led to altered expression of autophagy genes, but also affected other essential pathways for cell migration, proliferation, and homeostasis. The highly intertwined gene regulatory pathways show that in the cellular context, perturbation of one gene PC4 (Sub1 denoted as a green node) led to a disarrayed transcriptional output. Besides the autophagy pathway, the alterations of several of these autophagy related genes were linked to other cellular processes, the characteristics of which was observed in the PC4 depleted cells. As for example, ULK1 is critical for functioning of several cellular processes besides the autophagy machinery as mentioned like in cell motion, cell projection, etc. (Fig. 6C). Similarly, PTEN was found to regulate cell matrix adhesion, regulation of cell proliferation etc. Thus, perturbation of a single gene by the downregulation of the single nodal gene PC4, not only alters a single molecular pathway in the cells but also affects other interconnected cellular pathways. These highly concerted changes in the cells help us to understand better the altered characteristics of the PC4 depleted cells. Conclusively, alteration of an important nuclear protein PC4 led to an altered cellular state which might be a result of the combinatorial alteration of several genes which regulates several essential pathways for the proper maintenance and homeostasis of the cell.

## Discussion

Human transcriptional coactivator, PC4 has been previously reported to be an integral component of the chromatin.^12^ Depletion of PC4 at cellular level alters global organization of the nucleus as well as leads to an open chromatin state establishing it as one of the most important chromatin architectural protein. The chromatin architecture is known to be intricately modulated by the physiological state of the cell, a well-organized chromatin in a compact nucleus is essential for maintaining cellular homeostasis. Here we establish a novel role of the chromatin associated protein PC4 in cell segregation, chromosomal morphology, and maintenance of the epigenetic state of the cell. PC4 depleted cells show abnormal cellular segregation, enhanced hyper acetylation of histones and distorted shaped chromosomes. Interestingly, despite harbouring severe cellular defects, PC4 depleted cells shown higher proliferation rate than normal cells, and were also resistant to genotoxic stress like gamma irradiation, quite similar to the phenotype of an oncogenic transformed cells. To delineate the molecular mechanism of such a phenotype, we found that PC4 is a novel regulator of the well conserved cellular process of self-eating, called autophagy.

Autophagy levels were found to be highly elevated in absence of PC4. Electron micrographs of PC4 knockdown cells showed presence of various double membraned autophagy like vacuoles. Most of these vacuoles were found in close proximity to the irregularly shaped nucleus in PC4 knockdown cells. It was also interesting that PC4 depleted cells showed appearance of multivesicular bodies (large number of vacuoles engulfed in one vesicle). Appearance of multivesicular bodies has been reported in a particular type of autophagy process.^29^ Autophagy is a degradative pathway which plays an important role in maintaining protein homeostasis (proteostasis) as well as helps in the preservation of proper organelle function by selective removal of damaged organelles. The protective role of autophagy in maintaining cellular survival is mediated by the selective removal of dysfunctional mitochondria or other damaged organelles, which release pro-apoptotic factors and generate oxygen species.^30^ During autophagy, portions of cytosol are sequestered inside double-membraned vesicles (autophagosomes) that then fuse either with endocytic vesicles (as Multivesicular bodies) or lysosomes, which provide the hydrolytic enzymes that will degrade the content.^28^ The appearance of these multivesicular bodies and other double membraned vesicles especially near the nucleus in PC4 depleted cells thus might be possibly an indication of a cellular cue which might be operating via the altered nuclear architecture to a cytosolic process like autophagy. The enhanced autophagy levels in the PC4 knockdown cells play a critical role in its survivability possibly by maintaining cellular homeostasis in an otherwise physiological chaotic state.^18,30^

When the autophagy levels were depleted by administration of small molecule inhibitors or by knockdown of an essential autophagy gene ULK1, it highly compromised the survival rate of PC4 knockdown cells and upon gamma irradiation. This establishes the significant role of autophagy in attributing the property of enhanced growth rate to PC4 knockdown cells. Thus, PC4 depletion not only alters the nuclear architecture, chromatin compaction, epigenetic landscape but it also perturbs an important cytosolic event which in turn provides advantageous property to the otherwise abnormally transformed cells. We further investigated whether this phenomenon of enhanced autophagy is more specific to a loss of a chromatin condensing protein, PC4. PC4 knockdown significantly upregulated several autophagy related genes, like the AMPK and ULK genes which were previously reported to be prime regulators of the autophagy pathway. AMP-activated protein kinase (AMPK) is a well conserved energy sensing serine threonine kinase which is activated upon energy depleted or upon cellular stress conditions. This kinase further activates a cascade of proteins including the Unc-51-like kinase 1 and 2 (ULK1 and ULK2) which is an essential component in the formation and maturation of autophagosomes.^31^ We also find upregulation of MAP1LC3B gene in absence of PC4 which is involved in the final step of autophagosomes maturation and thereby degradation. This signifies that PC4 knockdown in cells not only leads to just an enhanced initiation of the self-eating process but the degradative pathway is complete and functional up to its final step. When we complemented PC4 in the knockdown background, we found significant downregulation of LC3 at protein levels (Fig. S2C and D) signifying its more direct role in regulation of autophagy.

We hypothesized that the phenomenon of enhanced autophagy in absence of PC4 might be occurring through the altered epigenetic state in PC4 depleted cells. The altered transcriptome in the PC4 depleted cells (Fig. S2A) might be due to the modified epigenetic landscape which is now permissive to enhanced transcription. The role of histone modifications in autophagy has not been deciphered in great details, however recent advances in the field, signify the role of the HDAC inhibitors like butyrate and suberoylanilide hydroxamic acid (SAHA) which enhanced autophagic cell death in various human cancer cell lines. Thus, there is increasing interest to look into histone hyperacetylation state and induction of autophagy.^32^ A very recent report suggests that the removal of H3K9me2 from the promoters of several autophagy-related genes resulted in an increase in autophagy levels in cells.^33^ Demethylation at the promoters of autophagy related genes at the time of autophagy induction is also known to occur, when the transcriptional repressor EHMT2 leaves the promoters resulting in subsequent acetylation of H3K9, thereby increasing the expression of autophagy-related proteins at the transcriptional level.^34^ Here in this report we try to directly co relate the histone acetylation level to autophagy induction in absence of PC4. PC4 depletion resulted in increased levels of histone modification marks related to transcriptional activation, like H3K9Ac, H3K27Ac, etc. We here show that PC4 knockdown enhanced the enrichment of the activation histone mark H3K9Ac mark at the promoters of AMPK and ULK1 and ULK2 genes, which possibly leads to an enhanced autophagy process. Thus, PC4 knockdown epigenetically upregulates the autophagy process by directly enhancing the autophagy related genes at the transcript level. This study provides one of the first direct evidences of interlink between chromatin, nuclear architecture and the process of autophagy.

Perturbation of the nuclear architecture by depletion of an important chromatin architectural protein, PC4, not only alters the nuclear state of the cells but also sends cues which regulates very important cytosolic events, like autophagy, to maintain cellular survivability under physiological stress. This observation has huge implications in understanding the very important roles of chromatin architectural protein in regulating the chromatin state and thereby regulating cytosolic processes which confer abnormal surviving ability of the cell particularly under pathophysiological conditions. However, whether the regulation of autophagy by PC4 is entirely due to the alteration of the epigenetic state of the cell or due to a response to cellular stress caused by the abnormal alteration of the nuclear state remains to be elucidated further. This study sheds a light on the link between chromatin architectural protein PC4 and its novel role in regulating autophagy to provide better survival property to the cells (Fig. 7). Enhanced autophagy in many cancer types empowers the cells to survive hypoxic and nutrient deprived environments.^35^ Histone modification profiles are altered in cancer cells as compared to normal cells.^36^ Emerging studies point at the histone modifications that accompany cancer and autophagy induction. Investigating the links between these three entities by employing cell lines such as sh-PC4 may shed light on these intricate cellular connections.

**Figure 7.**
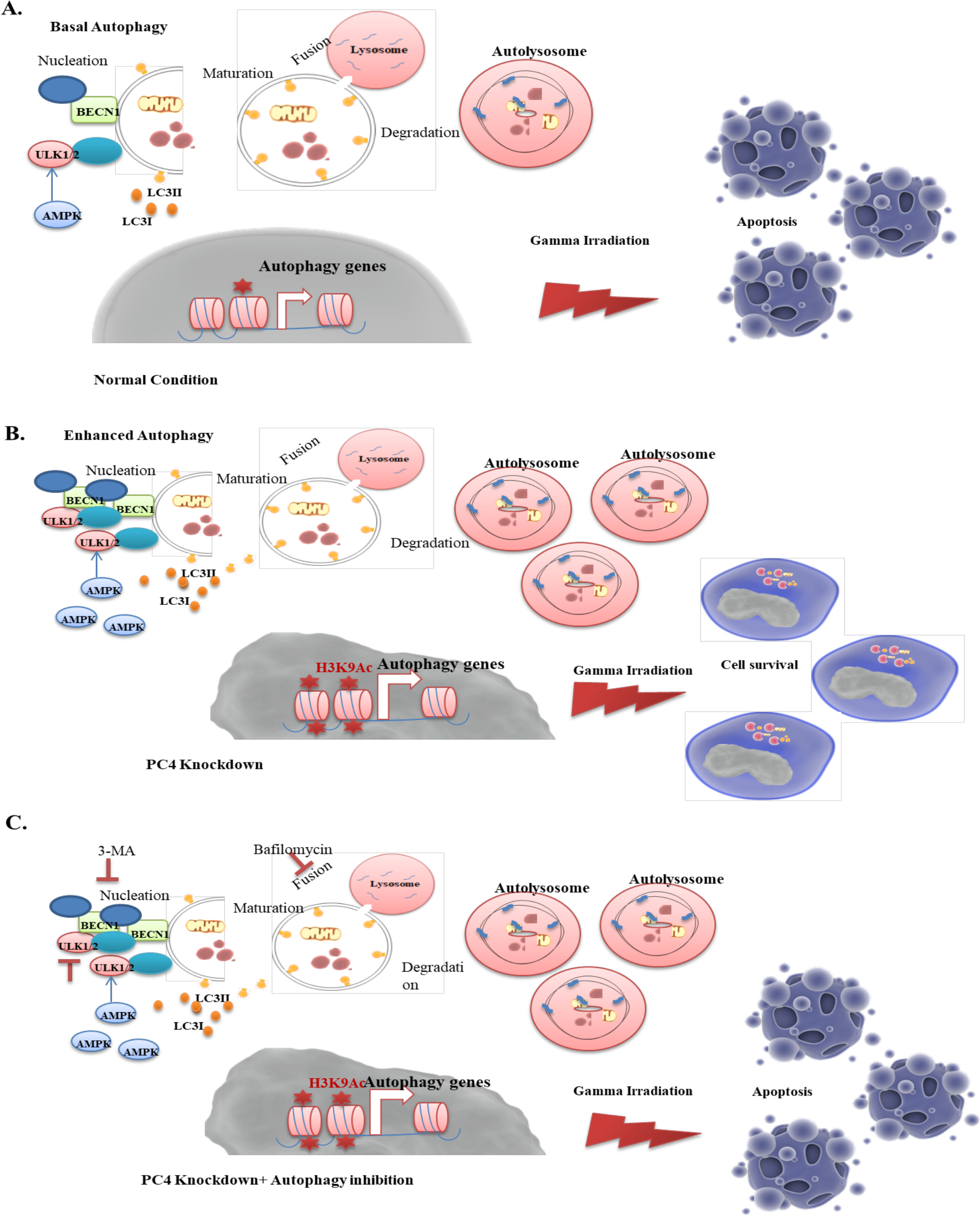
Model depicting the role of PC4 in autophagy regulation. (A) Control cells with basal autophagy undergo apoptosis upon gamma irradiation. (B) Enhanced autophagy in PC4 knockdown cells renders gamma irradiation resistance to these cells. (C) Genetic and chemical inhibition of autophagy makes the PC4 depleted cells susceptible to gamma irradiation mediated cell death.The figures have used illustration templates from the website somersault1824(http://www.somersault1824.com) available under a Creative Commons Attribution-Noncommercial-Share Alike License (CC BY-NC-SA 4.0)

## Materials and Methods

### Reagents and Media

Dulbecco’s Modified Eagle’s Medium (with 4500 mg/L glucose, L-glutamine, and 25 mM HEPES) (Sigma D1152), sodium bicarbonate (Sigma S5761), Penicillin-Streptomycin-Amphotericin B solution (Himedia, A002A), Foetal Bovine Serum heat inactivated (Life Technologies 04-121-1A). 3-Methyl adenine (Sigma 59281), Bafilomycin (Cayman Chemical, 11038), Earle’s Balanced Salt Solution (Sigma, E7510).

### Cell culture and treatment

Embryonic kidney cell line HEK293, HEK293T and HEK293 sh-PC4 were grown in the Dulbecco’s Modified Eagle’s Medium supplemented with sodium bicarbonate, 100 U/ml Penicillin 0.1 mg/ml Streptomycin, 0.25 μg/ml Amphotericin B and 10% heat inactivated foetal Bovine Serum at 37°C in 80% humidifier air and 5% CO_2_. For the starvation experiments, cells were allowed to grow upto 70% confluency and then were shifted to a low nutrient media of Earle’s Balanced Salt solution for 2 hours. For the small molecule inhibition assays the cells were grown to 70% confluency and then treated with different concentrations of 3-Methyl Adenine and Bafilomycin for 2 hours.

### PC4 knockdown stable cell line

This was generated using 10 μg pGIPZ lentiviral shRNAs targeting PC4 (Open Biosystems) and helper plasmids (5 μg psPAX2, 1.5 μg pRSV-Rev, 3.5 μg pCMV-VSV-G). 10 μg of sh-plasmid was mixed with helper plasmids (5 μg psPAX2, 1.5 μg pRSV-Revs, 3.5 μg pCMV-VSV-G) and were co transfected into HEK293T cells using calcium phosphate method. 48 hours post transfection media containing assembled virus was collected and its titre was estimated. Desired cell line (here HEK293) was infected with 10^5^ IU/ml virus. Infected cells were subjected to selection pressure 72 hours post transfection. Cells were grown in presence of 3 μg/ml puromycin for three passages to establish the cell line.

### Antibodies

PC4 (Laboratory reagent), histone H3 (Laboratory reagent), H3K9ac (Abcam, ab16635), H2AK5ac (Cell Signalling, 2576S), H4K16ac (Abcam, ab109463), Tubulin (Calbiochem, Merck DM1A) and GAPDH (Laboratory reagent), MAP1LC3B (Sigma L7543), H3K27Ac (Laboratory reagent), H4K12Ac (Laboratory reagent), ULK1 (Cell Signaling Technology, #8054).

### Gamma Irradiation Experiment

HEK293 and HEK293 sh-PC4 cells were seeded onto a 35mm dish and grown till 80% confluency. The dishes were exposed to different doses of radiation by setting the time in the irradiator (Blood Irradator-2000, Bl-2000), JNCASR, which has the rate of irradiation as 5.463 Grey units (Gy) per minute as follows:1 Gy-14 seconds;2 Gy-29 seconds;3 Gy-43 seconds. After exposure to radiation, fresh media was given to the cells and incubated for 24 hours at 37° C in a 5% CO_2_ supply and an 80% relative humidity of a CO_2_ incubator. After 24 hours of irradiation the cells were harvested and lysates were prepared for western blotting.

### Clonogenic assay

Clonogenic assay was performed using 200 cells (single cell population) seeded in 100 mm tissue culture dishes 24 hours prior to gamma irradiation. Cells were subjected to different doses of irradiation namely 0, 1, 2, 3 Gy. Media was replaced with fresh complete media soon after the irradiation. Plate was kept in 37°C incubator with 5% CO_2_ for 10 days to monitor colony formation. Colonies were stained with Crystal violet solution, counted in both the cases and surviving fraction was calculated as (No. of colonies at a particular dose/No. of colonies at 0Gy)*100. For Clonogenic assay with autophagy inhibitors, cells were seeded onto a 35mm dish and grown till 70% confluency, cells were then pre-treated with autophagy inhibitors for 2 hours, prior to exposure to 2 Gy gamma irradiation. After exposure to radiation, fresh media was given to the cells and incubated for 24 hours at 37° C in a 5% CO_2_ supply and an 80% relative humidity of a CO_2_ incubator. Cells which were pre-treated with autophagy inhibitors were also supplemented with fresh media containing the desired concentration of inhibitors after exposure to gamma irradiation. After 24 hours of irradiation, cells were trypsinized and then seeded for colony formation.

### shRNA mediated knockdown of ULK1

A specific shRNA against human ULK1 gene (Addgene # 27633) (shRNA sequence: ACATCGAGAACGTCACCAAGT) was transfected in the control cells as well as sh-PC4 cells. After 48 hours of transfection, the cells were trypsinized and seeded for colony formation assay as described earlier. Lysates collected from the shRNA transfected cell lines were subjected to immunoblotting after 48 hours of transfection against a specific antibody for ULK1.

### Real time RT-PCR

RNA was extracted from approximately 2× 10^4^ cells. 0.5 μg of RNA was reverse transcribed by using MMLV reverse transcriptase (Sigma, M1302). Resulting cDNAs were quantified by real-time PCR using KAPA SYBR FAST Universal qPCR master mix (KAPA Biosystems, KK4601) on the Stepone Plus realtime PCR platform (Life Technologies). Amplification with specific gene primers was performed. The housekeeping gene, β-actin was used for normalization. The primers used are given in supplementary table S1. Relative expression was normalized to the endogenous control Actin using the 2–ΔΔCt method. Experiments were carried out in technical triplicates, and biological duplicates. Primers used are listed in the table below.

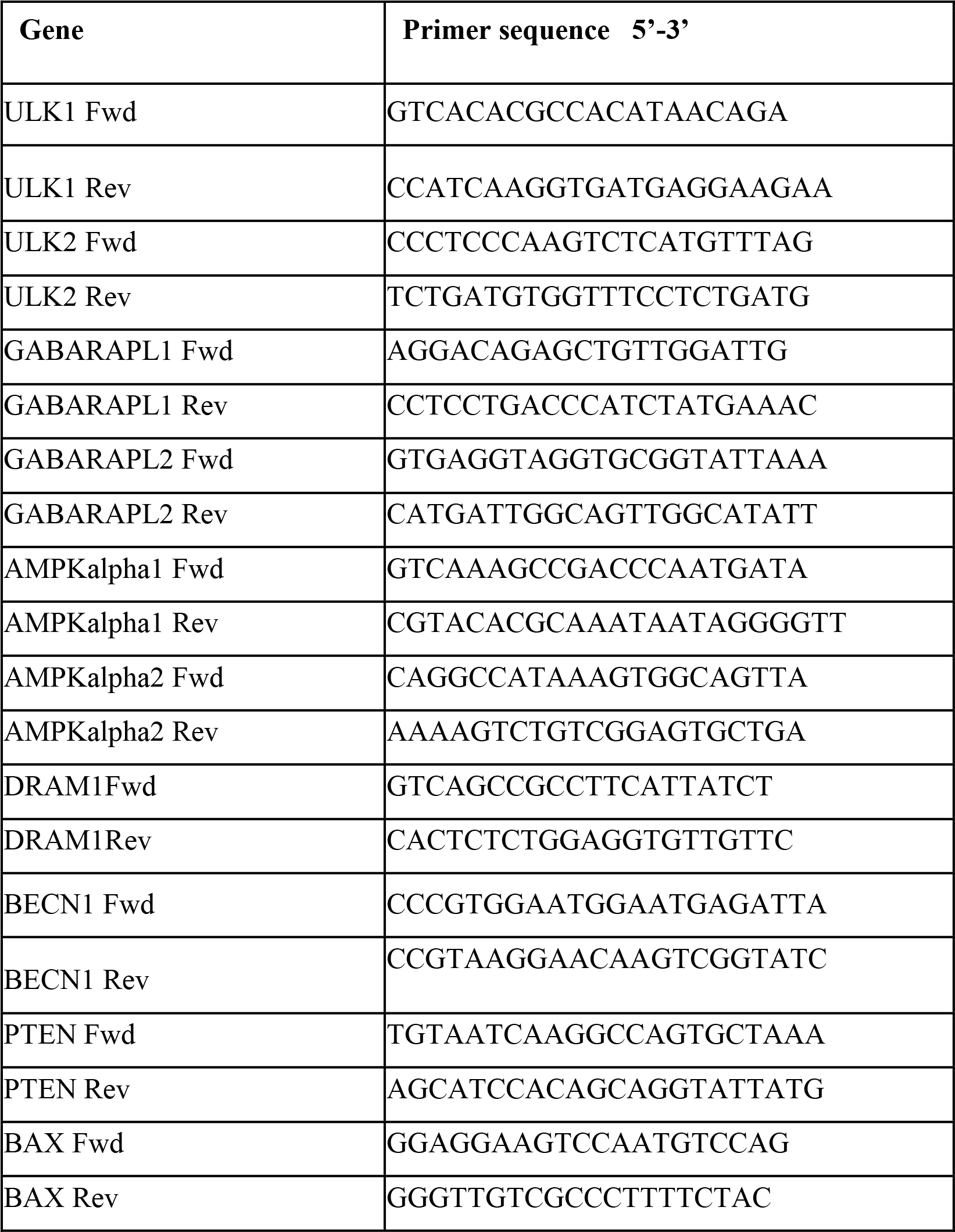

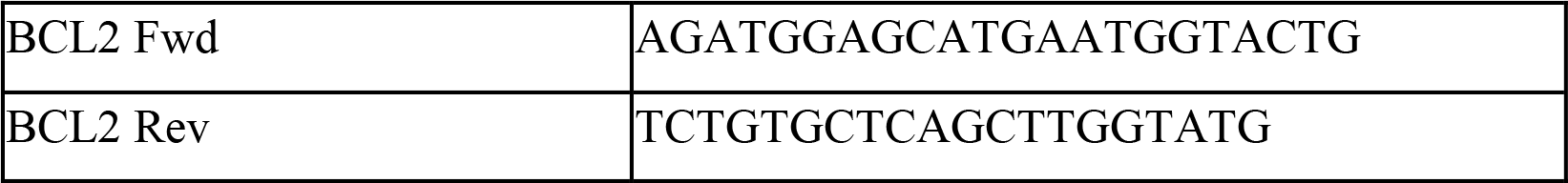

### Immunofluorescence

Assay was performed in desired cell lines and probing was carried out using different antibodies as indicated. DNA was stained with 0.1 μg /ml Hoechst 33258 (Sigma).

### Apoptosis Analysis and Cell sorting

Annexin V-Cy3 Apoptosis Detection Kit (Sigma APOAC) (utilizes the sharp and bright Cy3 red fluorescence dye conjugated to Annexin V. Annexin V-Cy3 binding can be detected by flow cytometry (Ex = 488 or 543 nm; Em = 570 nm) using the phycoerythrin emission signal detector (usually FL2). Detection of apoptosis was carried out as suggested by manufacturer’s kit protocol. Briefly, the cells were washed twice with Annexin Binding buffer IV (100 mM HEPES/NaOH, pH 7.5, containing 1.4 M NaCl and 25 mM CaCl_2_). Cells were dissolved in the buffer such that it is 10^6^ cells/ml. 100 μl of cell suspension was taken to this 1.5μl of Dye was added. Incubated for 15 mins in dark and then added to 400 μl of Annexin Buffer. This is used for analysis. For sorting 2 million cells were taken (100 ul suspensions containing 10^6^ cells stained with 1.5 μl of Dye). Volume was made upto 2 ml using Annexin Buffer. Cells which stained were excluded and the live population collected in autoclaved PBS buffer after sorting.

### Impedance measurement for proliferation/wound healing

The impedance across 8W10E array plates seeded with 10,000 cells per well was monitored in realtime using the ECIS-Zθ (Applied BioPhysics, USA) under multi-frequency mode. After saturation in the proliferation curve analyses were carried out using ECIS-Software-v1.2.8.PC and Excel 2007 (Microsoft) for the impedance values of grouped wells and plotted the average of triplicates for each population. At the end of saturation point wounds of equal size were created (at 5000 μA; 60000 Hz; 20 sec) and monitored the proliferation status over the next 12 hours. The normalized impedance was analyzed as described above.

### Scratch assay

This was performed using 80% confluent cells which were scratched with a sterile pipette tip in four separate places and image was captured after indicated time points.

### Cell lysate preparation

Cell pellets washed in 1X PBS from the desired cell line after the mentioned treatment were lysed in 1X SDS Buffer [50 mM Tris-Cl (pH 6.8),100 mM dithiothreitol, 2% SDS, 0.1% bromophenol blue, 10% glycerol]. Cell lysates were then heated at 90°C for 10 mins and then subjected to immunoblotting.

### Chromatin Immunoprecipitation

Chromatin immunoprecipitation was performed using HEK293 cells and HEK293 sh-PC4 cells, using the protocol described ^37^. The primers for the ChiP data were designed in accordance to the ENCODE data against the region in the promoters for each gene which showed a peak for H3K9Ac. The primers used for chiP qPCR are as follows: ULK1-Fw: 5′-GATTCCCAACCGGGATCAT-3′ and ULK1-Rev: 5′- CGTGCTTCTGAAAGCCAAAC-3′; ULK2-Fw: 5′- GCCAAGTTTCAGAACCACCTA-3′ and ULK2-Rev: 5′- GTGACTTCGAGTACAGCAAGAG-3′; AMPKα1-Fw: 5′-CTTTCCGACGACATGGTCTTTA-3′ and AMPKα1-Rev: 5′- CTGTAGGAGGCTGTCGATTTAC-3′; AMPKα2-Fw: 5′- CGCTGCACTGTGGGTAG-3′ and AMPKα2-Rev: 5′- CACGTAGTGTCCGATCTTCAC-3′; DRAM1-Fw: 5′- AGCGGTGGTGTAAAGTTGAT-3′ and DRAM1-Rev: 5′- GGTGCAGCTCCAGAGAATTTA-3′; HNF4a-Fw: 5′-TCCGTTGGCTCTGGATAATG-3′ and HNF4a-Rev: 5′- CACGTAGTGTCCGATCTTCAC-3′;

### Analysis of segregation defects

Wild type and PC4 knockdown HEK293 were grown to 60% confluence and treated with 26μg/ml Cytochalasin B for 3hours in 37°C incubator with 5% CO_2_. Cells were harvested by pipetting, washed once with 1X PBS and pellet was fixed in 1ml of Carnoy’s Fixative (Methanol: Glacial Acetic Acid 3:1). Fixed cells can be kept at 4°C for storage. Fixed cells were gently agitated and slides were prepared by the slide drop technique using Pasteur pipette. Slides were air dried, stained with 1mM of DAPI (4′,6-diamidino-2-phenylindole) and sealed with a coverslip. Imaging was carried out using Olympus BX-61 Fluorescence Microscope. Segregation defect was analyzed in terms of the number of anaphase bridges formed per 100 binucleates counted for both wild type and PC4 knockdown HEK293 and a graph was plotted using GraphPad Prism^TM^.

### FISH analysis of segregation defects

Wild type and PC4 knockdown HEK293 were grown to 60% confluence and both were treated with 26 μg/ml Cytochalasin B and 2.6% Colcemid for 3 hours in 37°C incubator with 5% CO_2_. Cells treated with Colcemid were harvested for chromosome analysis and treated with 1ml of 0.54% Potassium Chloride for 20 min at 37°C water bath. Pellet was then fixed in 1ml of Carnoy’s Fixative (Methanol: Glacial Acetic Acid 3:1) and stored at 4°C. Cytochalasin B treated cells for binucleate analysis were harvested, washed, fixed in 1ml of (Methanol: Glacial Acetic Acid 3:1) and stored at 4°C. Slides for chromosomes and binucleates were prepared by the slide drop technique, air dried and aged in a dust free environment overnight. Fluorescence In- Situ Hybridization of Telomere PNA probe was performed with chromosomes and binucleates as per the protocol^38^. Signal obtained was observed under Olympus BX 61 fluorescent microscope and quantified using ImageJ software^39^. A graph was plotted in GraphPad Prism^TM^ for the binucleate FISH in terms of the signal positivity in the daughter nuclei.

### Gene network analysis

For generating the core network representative of autophagy, significantly enriched biological categories / gene ontology / pathways harbouring differentially expressed genes upon treatment as compared to the control cells were subjected to network identification using Bridgen Island Software (Bionivid Technology Pvt. Ltd., Bangalore, India), resulting in identification of key nodes and edges. Output of Bridge Island Software was used as input to CytoScape V 2.8. Force directed spring embedded layout under yFiles algorithm was used to visualize the network that encompasses biological categories, differentially expressed genes that were significantly enriched. All the genes in the network were color coded based on their fold change upon PC4 knockdown, compared to control cells.

### Statistical analysis

All values are expressed as the mean ± S.E.M. Graphs were plotted in GraphPad Prism^TM^. For the statistical analysis, results were analysed using unpaired t test or one-way or two-way ANOVA and differences were considered significant if p< 0.01. All experiments were done in triplicates with biological replicates represented as n.

### Electron Microscopy

HEK293 and sh-PC4 cells were fixed in suspension with 4 % glutaraldehyde in 0.1 M cacodylate buffer (pH 7.3) after harvesting, overnight at 4°C. Cells were dehydrated with a graded series of ethanol, and embedded in epoxy resin. Then the areas containing cells were cut into ultrathin sections, stained and observed on transmission electron microscope Tecnai G2 F-30, a 300 Kv TEM / STEM equipped with a schottky field emission source and a point - point resolution of 2.2

## Acknowledgement

We acknowledge JNCASR, Department of Biotechnology, Government of India for the financial support. TKK is a recipient of Sir J.C. Bose National Fellowship. SS is a CSIR Senior Research Fellow. Part of this work was supported by Wellcome Trust/DBT India Alliance Intermediate Fellowship (500159-Z-09-Z) and JNCASR intramural funds to RM.

## List of abbreviations and acronyms

ULK1: Unc-51 Like Autophagy Activating Kinase 1
ULK2: Unc-51 Like Autophagy Activating Kinase 2
AMPKα1: 5’AMP-activated protein kinase catalytic subunit alpha-1
AMPKα2: 5’AMP-activated protein kinase catalytic subunit alpha-2
BECN1: Beclin1
DRAM1: DNA Damage Regulated Autophagy Modulator 1
PTEN: Phosphatidylinositol 3,4,5-trisphosphate 3-phosphatase and dual-specificity protein phosphatase
BCL2: Apoptosis regulator Bcl-2
3-MA: 3 Methyl adenine
shRNA: Short hairpin RNA
Baf: Bafilomycin A1
GABARAPL1: Gamma-aminobutyric acid receptor-associated protein-like 1
GABARAPL2: Gamma-aminobutyric acid receptor-associated protein-like 2
MAP1LC3B/LC3: Microtubule-associated protein 1B light chain 3B
MAP1LC3A: Microtubule-associated protein 1A light chain 3A

## References

1. Agresti A, Bianchi ME. HMGB proteins and gene expression. Curr Opin Genet Dev 2003; 13:170–178; PMID:12672494; https://doi.org/10.1016/S0959-437X(03)00023-6

2. Bustin M. Chromatin unfolding and activation by HMGN(*) chromosomal proteins. Trends Biochem Sci 2001; 26:431–437; PMID:11440855; https://doi.org/10.1016/S0968-0004(01)01855-2

3. Catez F, Yang H, Tracey KJ, Reeves R, Misteli T, Bustin M. Network of dynamic interactions between histone H1 and high-mobility group proteins in chromatin. Mol Cell Biol 2004; 24:4321–328; PMID:15121851; https://doi:10.1128/MCB.24.10.4321-4328.2004

4. Pallier C, Scaffidi P, Chopineau-Proust S, Agresti A, Nordmann P, Bianchi ME, Marechal V. Association of chromatin proteins high mobility group box (HMGB) 1 and HMGB2 with mitotic chromosomes. Mol Biol Cell 2003; 14:3414–3426; PMID:12925773; https://doi:10.1091/mbc.E02-09-0581

5. Li Y, Kirschmann DA, Wallrath LL. Does heterochromatin protein 1 always follow code? Proc Natl Acad Sci U S A 2002; 99:16462–16469; PMID:12151603; https://doi:10.1073/pnas.162371699

6. Kriaucionis S, Bird A. DNA methylation and Rett syndrome. Hum Mol Genet 2003; 12:R221–R227; PMID:12928486; https://doi:10.1093/hmg/ddg286

7. Kim MY, Mauro S, Gévry N, Lis JT, Kraus WL. NAD+-dependent modulation of chromatin structure and transcription by nucleosome binding properties of PARP-1. Cell 2004; 119:803–814; PMID:15607977; https://doi.org/10.1016/j.cell.2004.11.002

8. Das C, Hizume K, Batta K, Kumar BR, Gadad SS, Ganguly S, Lorain S, Verreault A, Sadhale PP, Takeyasu K, Kundu TK. Transcriptional coactivator PC4, a chromatin-associated protein, induces chromatin condensation. Mol Cell Biol 2006; 26:8303–8315; PMID:16982701; https://doi:10.1128/MCB.00887-06

9. Ballard DW, Philbrick WM, Bothwell AL.Identification of a novel 9-kDa polypeptide from nuclear extracts. DNA binding properties, primary structure, and in vitro expression. J Biol Chem 1988; 263:8450–8457; PMID:3372536

10. Ge H, Roeder RG. Purification, cloning and characterization of a human coactivator, PC4 that mediates transcriptional activation of class II genes. Cell 1994; 78:513–523; PMID:806239; https://doi.org/10.1016/0092-8674(94)90428-6

11. Kaiser K, Stelzer G, Meisterernst M. The coactivator p15 (PC4) initiates transcriptional activation during TFIIA-TFIID-promoter complex formation. EMBO J 1995; 14:3520–3527; PMID:7628453

12. Wang Z, Roeder RG. Roeder. DNA topoisomerase I and PC4 can interact with human TFIIIC to promote both accurate termination and transcription reinitiation by RNA polymerase III. Mol Cell 1998; 1:749–57; PMID:9660958; https://doi.org/10.1016/S1097-2765(00)80074-X

13. Mortusewicz O, Evers B, Helleday T. PC4 promotes genome stability and DNA repair through binding of ssDNA at DNA damage sites. Oncogene 2016; 35:761–770; PMID:25961912; https://doi:10.1038/onc.2015.135

14. Pan ZQ, Ge H, Amin AA, Hurwitz J. Transcription-positive cofactor 4 forms complexes with HSSB (RPA) on single-stranded DNA and influences HSSB-dependent enzymatic synthesis of simian virus 40 DNA. J Biol Chem 1996; 271:22111–22116; PMID:8703021; https://doi:10.1074/jbc.271.36.22111

15. Das C, Gadad SS, Kundu TK. Human positive coactivator 4 controls heterochromatinization and silencing of neural gene expression by interacting with REST/NRSF and CoREST. J Mol Biol 2010; 397:1–12; PMID:20080105; https://doi.org/10.1016/j.jmb.2009.12.058

16. Swaminathan A, Delage H, Chatterjee S, Belgarbi-Dutron L, Cassel R, Martinez N, Cosquer B, Kumari S, Mongelard F, Lannes B, et al. Transcriptional Coactivator and Chromatin Protein PC4 Is Involved in Hippocampal Neurogenesis and Spatial Memory Extinction. J Biol Chem 2016; 291:20303–20314; PMID:27471272; https://doi:10.1074/jbc.M116.744169

17. Polo SE, Jackson SP. Dynamics of DNA damage response proteins at DNA breaks: a focus on protein modifications. Genes Dev 2011; 25:409–433; PMID:21363960; https://doi:10.1101/gad.202131117.

18. Gewirtz DA. The Autophagic Response to Radiation: Relevance for Radiation Sensitization in Cancer Therapy. Radiat Res 2014; 182:363–367; PMID:25184372; https://doi:10.1667/RR13774.1

19. Lomonaco SL, Finniss S, Xiang C, Decarvalho A, Umansky F, Kalkanis SN, Mikkelsen T, Brodie C. The induction of autophagy by gamma-radiation contributes to the radioresistance of glioma stem cells. Int J Cancer 2009; 125:717–722; PMID:19431142; https://doi:10.1002/ijc.24402

20. Zhuang W, Li B, Long L, Chen L, Huang Q, Liang Z. Induction of autophagy promotes differentiation of glioma-initiating cells and their radiosensitivity. Int J Cancer 2011; 129:2720–2731; PMID:21384342; https://doi:10.1002/ijc.25975

21. Lin SY, Li TY, Liu Q, Zhang C, Li X, Chen Y, Zhang SM, Lian G, Liu Q, Ruan K,et al. GSK3-TIP60-ULK1 signaling pathway links growth factor deprivation to autophagy. Science 2012; 336:477–4781; PMID:22539723; https://doi:10.1126/science.1217032

22. Lee IH, Finkel T. Regulation of autophagy by the p300 acetyltransferase. J Biol Chem 2009; 284:6322–6328; PMID:19124466; https://doi:10.1074/jbc.M807135200

23. Dou Z, Xu C, Donahue G, Shimi T, Pan JA, Zhu J, Ivanov A, Capell BC, Drake AM, Shah PP, et al. Autophagy mediates degradation of nuclear lamina. Nature 2015; 527:105–109; PMID: 26524528; https://doi:10.1038/nature15548

24. Hewitt G, Korolchuk VI. Repair, Reuse, Recycle: The Expanding Role of Autophagy in Genome Maintenance. Trends Cell Biol 2017; 27:340–351; PMID:28011061; https://doi:10.1016/j.tcb.2016.11.011

25. Shin HJ, Kim H, Oh S, Lee JG, Kee M, Ko HJ, Kweon MN, Won KJ, Baek SH. AMPK–SKP2–CARM1 signalling cascade in transcriptional regulation of autophagy. Nature 2016; 534:553–557; PMID:27309807; https://doi:10.1038/nature18014

26. Glick D, Barth S, Macleod KF. Autophagy: cellular and molecular mechanisms. J Pathol 2010; 221:3–12; PMID:20225336; https://doi:10.1002/path.2697

27. Tanida I, Ikeguchi NM, Ueno T, Kominami E. Lysosomal Turnover, but Not a Cellular Level, of Endogenous LC3 is a Marker for Autophagy. Autophagy 2005; 1:84–91; PMID:16874052; https://doi:10.4161/auto.1.2.1697

28. Gozuacik D, Kimchi A. Autophagy as a cell death and tumor suppressor mechanism. Oncogene 2004; 23:2891–2906; PMID:15077152; https://doi:10.1038/sj.onc.1207521

29. Mizushima N, Levine B, Cuervo AM, Klionsky DJ. Autophagy fights disease through cellular self-digestion. Nature 2008; 451:1069–1075; PMID:18305538; https://doi:10.1038/nature06639

30. Vessoni AT, Filippi-Chiela EC, Menck CF, Lenz G. Autophagy and genomic integrity. Cell Death Differ 2013; 20:1444–1454; PMID: 23933813; https://doi:10.1038/cdd.2013.103

31. Shao Y, Gao Z, Marks PA, Jiang X. Apoptotic and autophagic cell death induced by histone deacetylase inhibitors. Proc Natl Acad Sci U S A 2004; 101:18030–18035; PMID:15596714; https://doi:10.1073/pnas.0408345102

32. Gammoh N, Marks PA, Jiang X. Curbing autophagy and histone deacetylases to kill cancer cells. Autophagy 2012; 8:1521–1522; PMID:22894919; http://dx.doi.org/10.4161/auto.21151

33. Artal-Martinez de Narvajas A, Gomez TS, Zhang JS, Mann AO, Taoda Y, Gorman JA, Herreros- Villanueva M, Gress TM, Ellenrieder V, Bujanda L. Epigenetic regulation of autophagy by the methyltransferase G9a. Mol Cell Biol 2013; 33:3983–993; PMID:23918802; https://doi:10.1128/MCB.00813-13

34. Füllgrabe J, Heldring N, Hermanson O, Joseph B. Cracking the survival code: autophagy-related histone modifications. Autophagy 2014; 10:556–561; PMID:24429873; https://doi:10.4161/auto.27280

35. White E. Deconvoluting the context-dependent role for autophagy in cancer. Nat Rev Cancer 2012; 12:401–410; PMID:22534666; https://doi:10.1038/nrc3262

36. Sharma S, Kelly TK, Jones PA. Epigenetics in cancer. Carcinogeneis 2010; 31:27–36; PMID:19752007; https://doi.org/10.1093/carcin/bgp220

37. Carey MF, Peterson CL, Smale ST. Chromatin immunoprecipitation (ChIP). Cold Spring Harb Protoc 2009; 2009:pdb.prot5279; PMID:20147264; https://doi:10.1101/pdb.prot5279

38. Poonepalli A, Banerjee B, Ramnarayanan K, Palanisamy N, Putti TC, Hande MP. Telomere-mediated genomic instability and the clinico-pathological parameters in breast cancer. Genes Chromosomes Cancer 2008; 47:1098–1099; PMID:18720522; http://dx.doi.org/10.1002/gcc.20608

39. Burgess A, Vigneron S, Brioudes E, Labbé J-C, Lorca T & Castro A. Loss of human Greatwall results in G2 arrest and multiple mitotic defects due to deregulation of the cyclin B-Cdc2/PP2A balance. Proc Natl Acad Sci U S A 2010; 107:12564–12569; PMID:20538976; https://doi:10.1073/pnas.0914191107

